# Museomics of a rare taxon: placing Whalleyanidae in the Lepidoptera Tree of Life

**DOI:** 10.1101/2020.08.18.255182

**Authors:** Victoria G. Twort, Joël Minet, Christopher W. Wheat, Niklas Wahlberg

**Affiliations:** Department of Biology, Lund University, Lund, Sweden; Muséum National d’Histoire Naturelle, ISYEB, Paris, France; Department of Zoology, Stockholm University, Stockholm, Sweden; The Finnish Museum of Natural History Luomus, Zoology Unit, University of Helsinki, Helsinki, Finland

## Abstract

Museomics is a valuable tool that utilises the diverse biobanks that are natural history museums. The ability to sequence genomes from old specimens has expanded not only the variety of interesting taxa available to study but also the scope of questions that can be investigated in order to further knowledge about biodiversity. Here we present whole genome sequencing results from the enigmatic genus *Whalleyana*, as well as the families Callidulidae and Hyblaeidae. Library preparation was carried out on four museum specimens and one existing DNA extract and sequenced with Illumina short reads. *De novo* assembly resulted in highly fragmented genomes with the N50 ranging from 317 – 2,078 bp. Mining of a manually curated gene set of 332 genes from these draft genomes had an overall gene recovery rate of 64 – 90%. Phylogenetic analysis places *Whalleyana* as sister to Callidulidae, and *Hyblaea* as sister to Pyraloidea. Since the former sister-group relationship turns out to be also supported by ten morphological synapomorphies, we propose to formally assign the Whalleyanidae to the superfamily Calliduloidea. These results highlight the usefulness of not only museum specimens, but also existing DNA extracts, for whole genome sequencing and gene mining for phylogenomic studies.

## Introduction

Natural history museums represent a diverse biobank of many interesting extant, rare and extinct taxa, making them an important scientific resource (Yeates et al. 2016; Graham et al. 2004; Suarez & Tsutsui 2004; Shaffer et al. 1998). Many species present in these collections are more accessible than in their original habitats due to a variety of factors (Thessen et al. 2012; Wandeler et al. 2007), such as remote geographical distributions, rare or endangered taxa, taxa that have since gone extinct, or taxa that have not been seen again following their initial collection. Although natural history collections have primarily been used for traditional morphological and taxonomic studies, ongoing advances in DNA sequencing technologies has expanded their role into the realm of genetics, and many other fields (Paijmans et al. 2013; Wiley et al. 2013; Shaffer et al. 1998). Despite the fact that these collections are now being recognized as an important genetic resource, most of the specimens in museums were collected prior to the use of DNA sequencing technology and were thus not preserved with the conservation of DNA material in mind. Hence, the DNA from these samples is often damaged and degraded and they were considered unsuitable for traditional molecular methods (Wandeler et al. 2007). In spite of this, museum specimens have been, and continue to be used for molecular studies (for example: Bi et al., 2013; Besnard et al., 2014, 2016; Chang et al., 2017). Initial studies focused on using PCR and Sanger sequencing of short fragments of genes (Cooper et al. 2006; Thomas et al. 1990; Houde & Braun 1988), however this approach not only requires the development of very specific primers for each gene, hence relying on prior genetic knowledge, but can also be cost prohibitive, and laborious (Wandeler et al. 2007; Soltis & Soltis 1993).

The development of high-throughput sequencing (HTS) technologies offers a promise of more efficient ways of sequencing DNA from museum specimens (Hofreiter et al. 2015; Rizzi et al. 2012). This is because HTS involves the sequencing of short fragments of DNA, which is a typical characteristic of DNA extracted from museum specimens, and results in large volumes of sequence data from relatively small amounts of starting material that provides good genome wide genetic data. The publication of the mammoth genome (Poinar et al. 2006), followed by the Neanderthal genome (Meyer et al. 2012; Green et al. 2010) showed the promise of HTS technologies and ancient DNA. Since then, HTS has slowly been applied to a more diverse range of taxa. One particular application being widely used is targeted sequence capture, in which focal regions of the genome are isolated and sequenced. The regions targeted are typically those that are well conserved across the taxa of interest, such as exons and ultraconserved elements (UCEs), so that only a single probe set is required to be designed for the group(s) of interest. These target-based approaches have been applied to a variety of taxa and specimen ages (for example: Bi et al., 2013; Staats et al., 2013; Bailey et al., 2016; Blaimer, Lloyd, Guillory, & Brady, 2016; Prosser, Dewaard, Miller, & Hebert, 2016), and have proven relatively successful in recovering the regions of interest for phylogenomic studies. Despite these advances, application of these methods requires prior genetic knowledge of the taxa of interest in order to facilitate probe design.

An alternative approach to targeted sequence capture is whole-genome sequencing. Although initial studies focused on taxa for which a reference genome, or one closely related, already existed (for example: Rowe et al., 2011; Staats et al., 2013), *de novo* based approaches are starting to be used. Nevertheless, these studies tend to be carried out with taxa for which large volumes of starting material are available for DNA extraction, with very few studies using HTS based approaches on old insect specimens (Kanda et al. 2015; Heintzman et al. 2014; Maddison & Cooper 2014; Tin et al. 2014; Staats et al. 2013). Initial studies focused on recovering specific regions, such as mitochondrial or ribosomal DNA, which are present in multiple copies per cell (Heintzman et al. 2014; Staats et al. 2013). Kanda *et al.* (2015) highlighted that the recovery of low-copy regions of the genome is possible from a diversity of museum specimens spanning a variety of ages, preservation methods and DNA quality. The use of both *de novo* and reference-based assemblies for gene recovery highlighted that although more loci can be recovered if one has an existing reference, many loci are still obtained with a *de novo* approach (Kanda et al. 2015). Additionally, Sproul and Maddison (2017) presented a non-destructive DNA extraction method for historical beetle samples and successfully prepared and sequenced libraries from the low amounts of degraded DNA recovered with good results. These studies combined show that for historical samples of interest it is possible to get enough DNA for sequencing and gene recovery without the need for any prior genetic knowledge.

Here we explore the application of museomics to resolve the phylogenetic position of the enigmatic *Whalleyana* moths. The genus *Whalleyana* is endemic to Madagascar, and was first described in 1977 as an odd member of the Thyrididae (Viette 1977). Very little is known about the two species (*Whalleyana vroni* Viette, 1977 and *W. toni* Viette, 1977) that make up this genus, including their phylogenetic placement within Lepidoptera. In addition to the lack of knowledge, neither species has been collected since the 1990’s (David C. Lees, pers. comm. to JM). Thus, in order to get a better understanding of this taxon, existing museum samples are the only resource available. The aim of this study is to utilize low-coverage whole genome sequencing of interesting museum specimens and existing DNA extracts to highlight the usefulness of such approaches for answering questions, such as where in the Lepidoptera phylogeny does *Whalleyana* belong. We also sequence museum specimens or existing genomic DNA extracts of potentially related taxa belonging to the families Callidulidae and Hyblaeidae. Such approaches not only allow us access to taxa we may not have had the opportunity to investigate previously, but by utilizing a whole genome sequencing approach we are able to generate a rich dataset for future researchers using diverse approaches. We assessed the robustness of our results with morphological data mainly from the adult stage. Indeed, the early stages of the Whalleyanidae remain completely unknown.

## Material and Methods

### Taxon sampling for molecular analyses

Whole genome sequencing data were generated from museum specimens of the following species: *Whalleyana vroni* (collected in 1969), *Helicomitra pulchra* Butler, 1878 (collected in 1974), *Griveaudia vieui* Viette, 1958 (collected in 1969), *Hyblaea madagascariensis* Viette, 1961 (collected in 1975), as well as a DNA extract of *Hyblaea puera* (Cramer, 1777) from Mutanen et al. (2010). *Helicomitra* and *Griveaudia* belong to the family Callidulidae, and *Hyblaea* to Hyblaeidae. Both families are potentially related to *Whalleyana*, and the latter has no genomic resources available prior to our study. All museum specimens are from the Muséum National d’Histoire Naturelle, Paris (MNHN). The DNA extract of *H. puera* was kindly loaned by Marko Mutanen (University of Oulu, Finland).

### DNA extraction

DNA was extracted from the abdomens of four specimens, using the QIAamp DNA Micro kit (Qiagen) following the manufacturer’s protocol, with the following modifications: no crushing of the samples was carried out prior to incubation with lysis buffer, following overnight incubation samples were centrifuged and the supernatant carried forward for extraction with the remaining tissue being placed in ethanol, and elution buffer was incubated in the columns for 20 minutes at room temperature prior to elution. We took reasonable measures to avoid contamination, including the use of filter tips and sterilized work areas that are physically separated from areas where fresh specimens are prepared. The resulting DNA extracts were visualized on 0.8% w/v agarose gels stained with SYBR safe (Fisher Scientific) to determine DNA fragmentation levels. In the case of the *H. puera* extract, due to high molecular weight DNA, DNA was sonicated to approx. 200 – 300 bp fragments using a Bioryptor^®^ with the following settings: (M) medium power output, 30 sec ON/ 90 sec OFF pulses for 30 minutes in a 4°C water bath, followed by vacuum centrifugation and resuspension in 50 μl of elution buffer.

### Library Preparation and Sequencing

Library preparation followed a modified protocol of Meyer and Kircher (2010). All reagent distributors and catalogue numbers are given in Table S1. Firstly, DNA was blunt-end repaired. The reaction mix consisted of: 1x Tango Buffer, dNTP (100 μM each), 1 mM ATP, 0.5 U/μl PNK and 0.1 U/μl T4 DNA Polymerase. The reaction was incubated for 15 min at 25°C, followed by 5 min at 12°C. Purification of the reaction was carried out with the MinElute purification kit (Qiagen), and elution in 22 μl EB buffer. Adapter ligation followed purification with a reaction mix containing: 1x T4 ligation buffer, 5% PEG-4000, 0.125 U/μl T4 Ligase, and an adapter mix of P7 and P5 adapters (Meyer & Kircher 2010) 2.5 μM each. Reactions were incubated for 30 min at 22°C. After purification with the MinElute purification kit (Qiagen), adapter fill-in was performed using the following reaction mix: 1x Isothermopol amplification buffer, dNTP (250 μM) and 0.3 U/μl Bst polymerase. Incubation at 37°C for 20 min, was followed by the final heatkill was performed by incubation for 20 min at 80°C.

Indexing and amplification of each library was carried out with 3 μl of library template and a unique dual indexing strategy. The amplification mix consisted of 0.05 U/μl AccuPrime Pfx DNA Polymerase, 2.5 μl AccuPrime reaction mix, 200 nM IS4 primer (Meyer & Kircher 2010) and 200 nM of indexing primer. Amplification was carried out under the following conditions: 95°C for 2 min, 18 cycles of: 95°C for 15 s, 60°C for 30 s 68°C for 60 s, which were carried out in six independent reactions, to avoid amplification bias, and pooled prior to purification. Purification along with size selection was carried out using a two-step process with Agencourt AMPure XP beads. An initial bead concentration of 0.5X was used to remove long fragments that are likely to represent contamination from fresh DNA, libraries were selected with a bead concentration of 1.8X to size select the expected library range of 100 – 300 bp. The resulting libraries were quantified and quality checked with Quanti-iT™ PicoGreen™ dsDNA assay and with a DNA chip on a Bioanalyzer 2100, respectively. Multiplexed libraries were pooled as follows: *W. vroni* was pooled at 50% molar concentration in a pool of 9 samples and sequenced over two runs, while the remaining specimens were pooled in equimolar concentrations in a pool of 6 samples and sequenced using the Illumina HiSeq2500 technology with 150 bp paired-end reads.

### Genome Assembly

Raw reads were quality checked with FASTQC v0.11.8 (Andrews 2010). Sequencing reads resulting from samples with highly degraded DNA were treated from this point as single end reads. This approach was chosen, as degraded DNA is likely to randomly ligate together during the adapter ligation stage of library preparation, resulting in chimeras of different genomic regions (Willerslev & Cooper 2005). Nevertheless the sequencing information contained in the reads is still reliable, as chimera formation typically results in DNA inserts larger than read length, therefore more reliable results are obtained by treating data as single-end (Rowe et al. 2011). For the sample that underwent sonication (*H. puera*), reads were carried forward as paired-end. Reads with ambiguous bases (N’s) were removed from the dataset using Prinseq 0.20.4 (Schmieder & Edwards 2011). Trimmomatic 0.38 (Bolger et al. 2014) was used to remove low quality bases from their beginning (LEADING:3) and end (TRAILING:3), by removing reads below 30 bp, and by evaluation read quality with a sliding window approach. Quality was measured for sliding windows of 4 base pairs and had to be greater than PRHED 25 on average. The resulting cleaned reads were used for *de novo* genome assembly with spAdes v3.13.0 (Bankevich et al. 2012) with kmer values of 21, 33 and 55. The completeness of each assembly was assessed using BUSCO 3.0.2b (Simão et al. 2015) using the Insecta lineage set. However, due to the fragmented nature of the genomes BUSCO has trouble identifying orthologs, therefore the genomes were searched for the insecta lineage set using a tblastn approach (e-value threshold 1e-5, minimum identity of 60%) with standalone BLAST 2.9.0 (Camacho et al. 2009).

### Orthologue Identification and Alignment

Orthologues were identified from the fragmented genome assemblies using MESPA v1.3 (Neethiraj et al. 2017), with a custom set of 332 representative gene markers (11 of which are mitochondrial), which have been manually vetted for alignment and orthology based on their amino acid sequences from a set of 200 taxa of Lepidoptera. The details of the vetting process are described in Rota et al. (in prep.). The resulting DNA sequences were aligned to pre-existing reference alignments (taken from Rota et al. in prep.) based on their translated amino acid sequences using MAFFT v7.310 (Katoh et al. 2002) using the add fragments and auto options which keeps existing gaps in the alignments and chooses the most appropriate alignment strategy. The resulting amino acid alignments were manually checked to ensure accuracy, screen for the presence of pseudogenes, reading frame errors and alignment errors using Geneious 11.0.3 (https://www.geneious.com). The amino acid alignments were then converted back to nucleotide alignments, and the aligned DNA sequences were curated and maintained using the Voseq database (Peña & Malm 2012), which allows users to custom-make datasets for downstream phylogenetic analyses in chosen formats (e.g. FASTA, Nexus or Phylip formats). Raw sequencing data can be found under Bioproject PRJNA631866, while genome assemblies can be accessed from Zenodo, DOI: 10.5281/zenodo.3629334

### Phylogenetic analysis

In order to investigate the phylogenetic placement of *Whalleyana*, the new sequences were added to a manually curated dataset derived from published transcriptomes and genomes of ditrysian Lepidoptera (Rota et al. in prep). The final dataset consisted of a total of 338 gene fragments, spanning 332 genes, across 169 taxa (164 taxa were taken from Rota et al. (in prep.), See Table S2 for full list of included taxa and Table S3 for the genes recovered from the specimens in this study), with the final alignment file being created in Voseq.

The resulting nucleotide and amino acid (aa) sequence alignments were analysed in a maximum likelihood framework using the program IQ-TREE (Nguyen et al. 2014). For the nucleotide dataset, third codon positions were removed (nt12). Nucleotide data were also analysed using degen1 coding (Zwick et al. 2012; Regier et al. 2010). Each dataset was analysed partitioned by gene, with ModelFinder (Kalyaanamoorthy et al. 2017) run first, and then the maximum likelihood search run after based on the optimal models found for each gene. The robustness of our phylogenetic hypotheses was assessed with 1000 ultrafast bootstrap (UFBoot2) approximations (Hoang et al. 2017) in IQ-TREE. Analyses were run on the CIPRES portal (Miller et al. 2010).

### Morphological analyses

Adult morphology was investigated using a collection of specimens dissected by one of us and belonging to the MNHN. These specimens had been chosen to represent most ditrysian families within the framework of several previous publications (e.g. Minet, 1991 and Rajaei et al., 2015). Among the families which are more precisely the focus of the present study, specimens that have been entirely dissected belong to the following genera: *Striglina* Guenée, 1877 (Thyrididae: Striglininae), *Marmax* Rafinesque, 1815 (Thyrididae: Charideinae), *Thyris* Laspeyres, 1803 (Thyridinae), *Chrysotypus* Butler, 1879 (Thyrididae: Siculodinae), *Rhodoneura* Guenée, 1858 (same subfamily), *Whalleyana* Viette, 1977 (Whalleyanidae), *Helicomitra* Butler, 1878 (Callidulidae: Pterothysaninae), *Griveaudia* Viette, 1958 (Callidulidae: Griveaudiinae), *Callidula* Hübner, 1819 (Callidulinae), and *Hyblaea* Fabricius, 1794 (Hyblaeidae). After removal of their wings, these imagos were macerated in a hot 10% potassium hydroxide solution (KOH), rinsed in demineralized water, then cleaned, descaled, stained (with Chlorazol Black E), and dissected in 70% ethanol (following methods expounded by Brock, 1971 and Robinson, 1976). Afterwards, the various parts of the body were severed from adjacent regions and either stored intact in 70% ethanol or preserved as permanent slide mounts in Euparal (following standard techniques: Robinson, 1976). Structures of possible phylogenetic interest were photographed and/or examined using an Olympus SZH stereo microscope with a linear magnification range of X7.5 to X128. In the search of apomorphic traits suited to support, or not, our molecular phylogeny, special attention was paid to the less homoplastic characters, nevertheless without neglecting any character easy to polarize through outgroup comparisons. External characters whose observation does not require dissections were surveyed on a large scale and full account was taken of published morphological data, especially in the case of the hyblaeoid family Prodidactidae (Kaila et al. 2013; Epstein & Brown 2003).

## Results

### a) The molecular approach

DNA extraction of the four museum specimens were successful with DNA fragments ranging from 70 - 300 bp in size, while sonication of the *H. puera* extract resulted in DNA fragment lengths of 300 bp (results not shown). Sequencing resulted in a total of 754 million reads across the runs, ca. 462 million reads belonged to *W. vroni*, and an average of 72 million reads for the other four samples (Table 1). Of these reads >86% passed adapter and quality trimming. Each sample was *de novo* assembled. The resulting genome assemblies were highly fragmented with average contig lengths of 321 bp and N50’s ranging from 317 – 2,078 bp (Table 1). Assessment of the completeness of the resulting assemblies with BUSCO, highlighted the difficulty the program has in finding ortholgues in fragmented genomes (results not shown). However, the ortholgue set can still be used with a BLAST approach to assess presence of the conserved genes. The blast search for the 1, insecta orthologues showed the majority of orthologues are present, in at least fragmented form with between 74% and 87% being present in the genomes (Table 1).

**Table 1:** Genome Assembly and Gene recovery statistics

| Sample | *Griveaudia vieui* | *Hyblaea madagascariensis* | *Helicomitra pulchra* | *Whalleyana vroni* | *Hyblaea purea* |
| --- | --- | --- | --- | --- | --- |
| Code | VT58 | VT57 | VT56 | VT11 | MM07227 |
| Data treated as | SE | SE | SE | SE | PE |
| Raw Reads (Paired) | 73,451,727 | 81,670,390 | 68,286,636 | 462,344,781 | 68,569,202 |
| Cleaned Read Pairs | --- | --- | --- | --- | 23863949 |
| Cleaned Read Unpaired R1 | 65,227,946 | 71,533,273 | 61,719,435 | 422,057,592 | 37,539,693 |
| Cleaned Reads Unpaired R2 | 63,764,833 | 69,860,500 | 58,962,343 | 407,003,611 | 391,589 |
| Contigs | 700,194 | 746,054 | 1,155,426 | 1,639,567 | 985,209 |
| Max Contig Length | 5,413 | 4,792 | 5,441 | 7,920 | 65,231 |
| Minimum Contig Length | 56 | 56 | 56 | 56 | 56 |
| Average Contig Length | 284.0 ± 147.9 | 330.2 ± 183.0 | 278.5 ± 197.7 | 295.5 ± 330.1 | 417.4 ± 1180.6 |
| Median Contig Length | 264.0 | 291.0 | 250.0 | 209.0 | 108.0 |
| Total Contig Length | 198,848,300 | 246,324,962 | 321,785,346 | 484,482,679 | 411,261,707 |
| % non ATCG characters | 0.001 | 0.001 | 0.005 | 0.01 | 0.287 |
| Contigs >= 100 bp | 610,227 | 683,788 | 941,766 | 1,104,568 | 545,487 |
| Contigs >= 200 bp | 577,430 | 653,684 | 735,144 | 835,486 | 301,242 |
| Contigs >= 500 bp | 51,490 | 101,611 | 135,739 | 266,889 | 139,271 |
| Contigs >= 1 Kbp | 1,429 | 6,611 | 8,871 | 65,308 | 84,638 |
| Contigs >= 10 Kbp | --- | --- | --- | --- | 2,639 |
| N50 value | 317 | 367 | 361 | 478 | 2,078 |
| **Numer of Genes Recovered (/338)** | 215 (63%) | 244 (72%) | 244 (72%) | 255 (75%) | 304 (90%) |
| **Number of BUSCO Insecta Genes Identified (/1,658)** | 1,201 (72%) | 1,415 (85%) | 1,396 (84%) | 1,435 (87%) | 1,400 (84%) |

Identification of the curated Lepidoptera gene set with MESPA had a recovery rate of between 64% −90% (Tables 1 and S3). The resulting sequences were uploaded to an in-house database (Voseq Peña & Malm, 2012), and a final concatenated dataset comprising a total of 162 taxa, and 291,516 nucleotides in length was used for analyses. Analysis of both the nucleotide and amino acid datasets shows stable placement of *Whalleyana* as sister to Callidulidae, and *Hyblaea* as sister to Pyraloidea (Fig. 1). Both of these relationships have 100% UFbootstrap support regardless of data form, except for nt12, where the sister relationship of *Hyblaea* and Pyraloidea receives only 86%.

**Figure 1.**
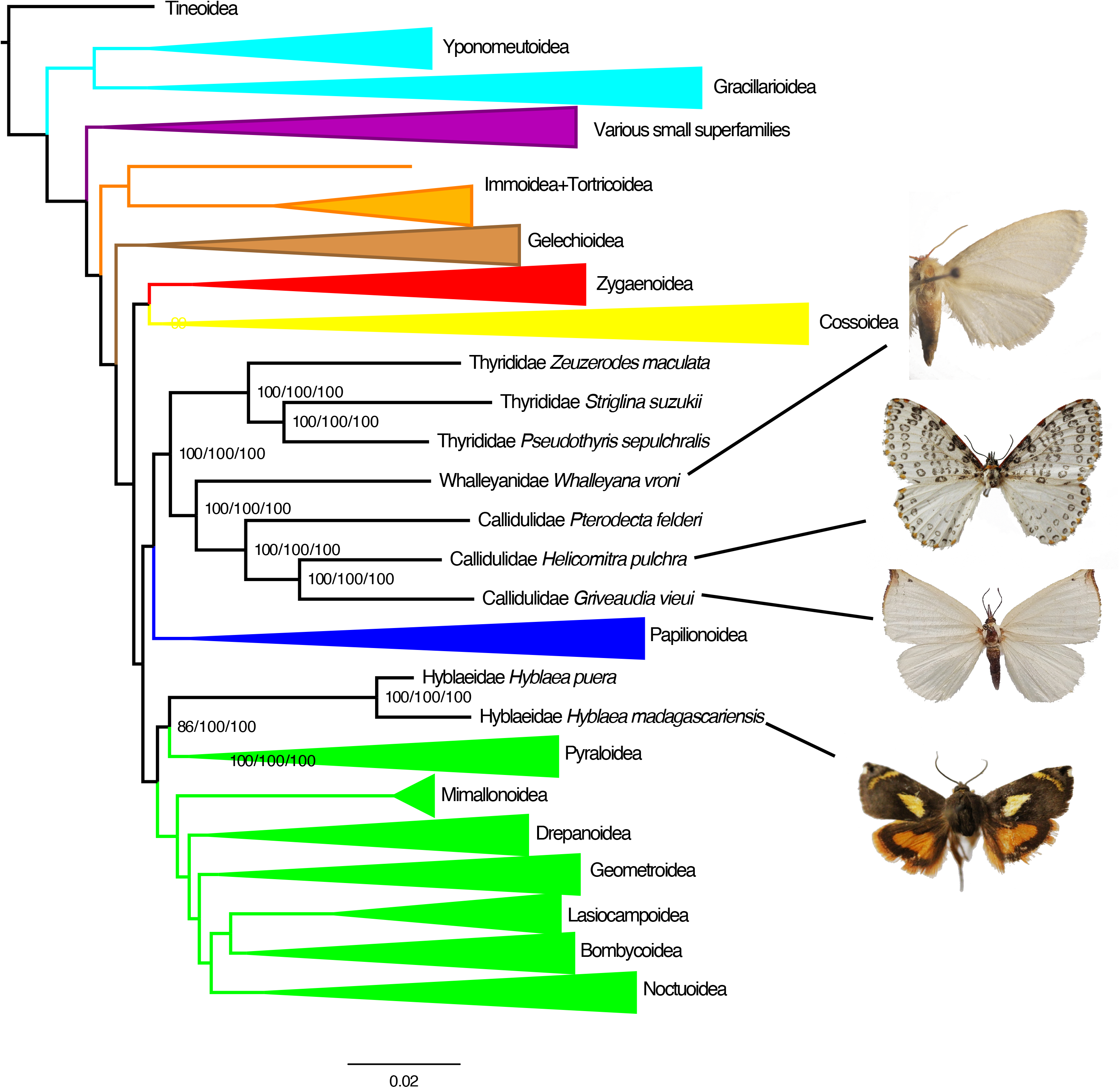
Phylogenetic relationships of *Whalleyana*, *Helicomitra*, *Griveaudia* and *Hyblaea* based on 332 genes. Superfamilies that are not relevant to these taxa are shown as collapsed. Numbers to the right of each node give the UltraFastBootstraps for each dataset analysed: nt12/degen1/aa.

### b) Morphological synapomorphies and autapomorphies

The interpretation of the following characters is based on Fig. 1, but takes also into account the uncertainties mentioned hereafter (see Discussion) about the interrelationships between Gelechioidea, Papilionoidea and the Thyridoidea + Calliduloidea lineage.

Thyrididae, Whalleyanidae and Callidulidae share six imaginal synapomorphies: (1) on the head, the ocellus is either absent or devoid of a distinct lens (as also in all Papilionoidea and Hyblaeoidea); (2) in the forewing, vein M2 arises closer to M3 than to M1 (a frequently encountered apomorphy, which also occurs in Hyblaeoidea + Pyraloidea but does not pertain to the ground plan of Papilionoidea nor to that of Gelechioidea: for instance, in the latter superfamily, M2 arises closer to M1 than to M3 in such genera as *Hypertropha* Meyrick, 1880 and *Donacostola* Meyrick, 1931); (3) at the base of the forewing, the spinarea is absent or extremely small (an apomorphy also present in all Papilionoidea but absent in the genus *Hyblaea* Fabricius, 1793, which retains a large spinarea (Common, 1990: Fig. 108), and in many Gelechioidea and Pyraloidea); (4) both fore- and hindwings lack a distinct, tubular CuP (this vein being replaced by a fold, which may resemble a vein in certain large Thyrididae (e.g. in the genus *Draconia* Hübner, 1820); by contrast a true vein CuP is preserved in both pairs of wings of Prodidactidae (Hyblaeoidea) (Epstein & Brown 2003: Fig. 9) and in the hindwing of *Hyblaea*; (5) in the hindwing, vein Sc + R is approximated to, or fused with, vein Rs (beyond wing base and either before or beyond the upper angle of the discal cell) (in the genus *Prodidactis* Meyrick, 1921, only the base of Sc is approximated to the upper edge of the hindwing discal cell); (6) in the male genitalia, the juxta is provided with a pair of erect “arms” that are directed caudad or dorsad (Fig. 2A, arrow; since Whalleyanidae and Callidulidae appear as sister groups, the absence of these erect arms in Callidulidae should represent a loss rather than a primary condition). A larval trait may represent a seventh synapomorphy of these families (when the larva of *Whalleyana* is discovered), namely the presence of just one seta in the L group of segment A9 (see e.g. Fig. 7 in Chistyakov et al. (1994)). While this apomorphy also occurs in *Hyblaea* and many Pyraloidea, two L setae are preserved (on A9, laterally) in Prodidactidae (Epstein & Brown 2003: Fig. 14) and three in the ground plan of the pyraloid larva (Neunzing 1987: 463). Thyrididae and Callidulinae also share the following pupal apomorphy: the mandibles (pilifers sensu Mosher 1916) are distinctly adjacent on the meson (Nakamura 2011: Figs 1, 2 and 5). Nevertheless, they are not adjacent in the subfamily Pterothysaninae of Callidulidae (Nakamura 2011: Fig.3) while the pupa of the Griveaudiinae remains unknown to date.

**Figure 2.**
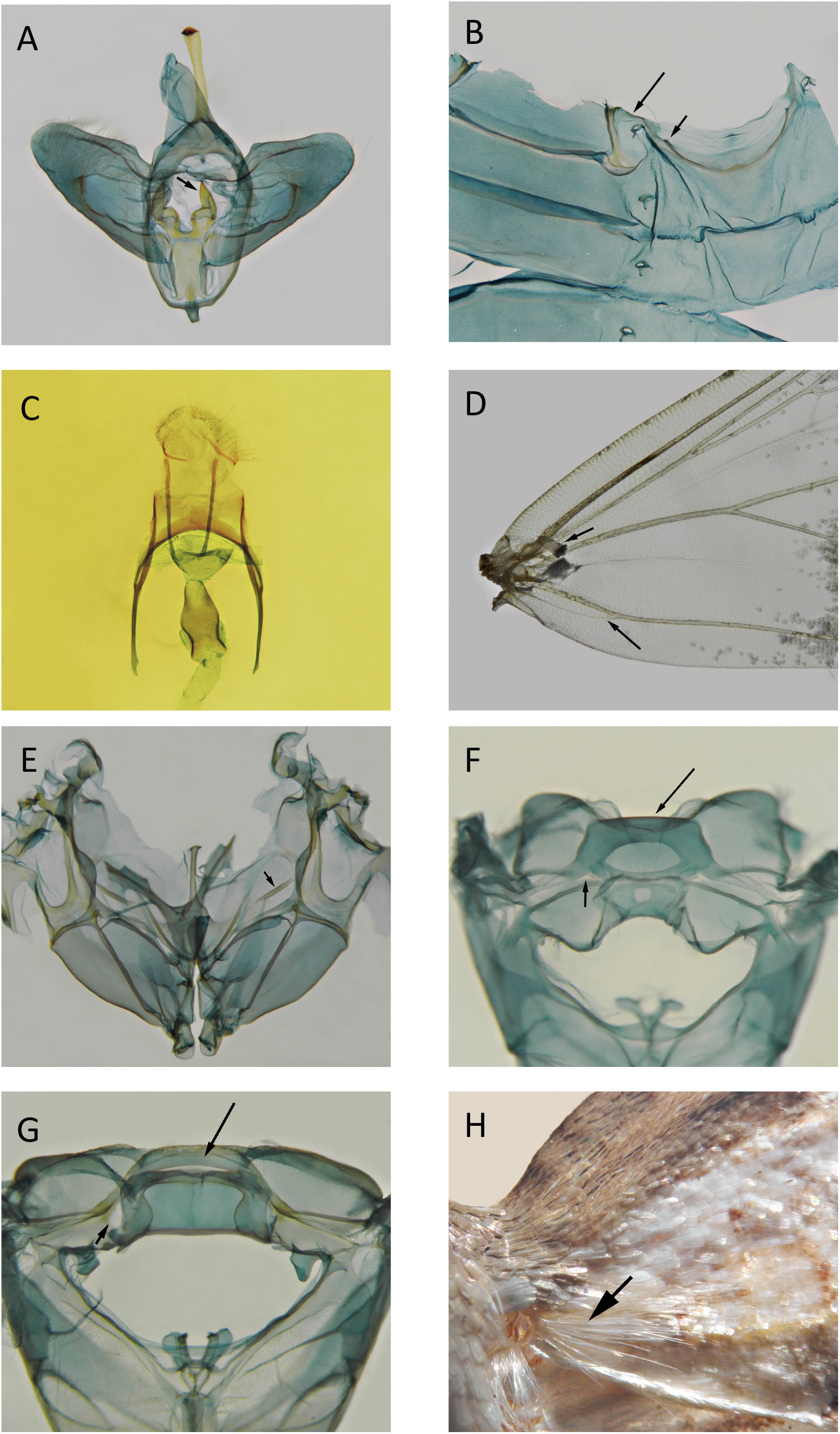
A. *Whalleyana vroni*, male genitalia (arrow: one of the two free, dorsally directed, arms of the juxta). B. *Whalleyana vroni*, first three segments of the male abdomen, with the sterna on the right (short arrow: apodeme; long arrow: tergosternal sclerite). C. *Whalleyana toni*, posterior region of the female genitalia (*preparation* P. Viette 5451). D. *Whalleyana vroni*, base of the male forewing (ventral surface) after removal of most scales (short arrow: retinaculum; long arrow: lower branch of the “anal fork”). E. *Whalleyana vroni*, mesothoracic pleurosternum in anterior view (arrow: precoxal sulcus). F. *Griveaudia vieui*, metathorax in posterior view (short arrow: left-hand fenestra lateralis; long arrow: scutellum). G. *Whalleyana toni*, metathorax in posterior view (short arrow: ventral edge of the left-hand fenestra lateralis; long arrow: scutellum). H. *Rhodoneura opalinula*, forewing base in dorsal view (arrow: bunch of piliform scales arising from the base of vein A1).

Ten synapomorphies from adult morphology clearly support a sister-group relationship of Whalleyanidae and Callidulidae: (7) in the antennae of dried specimens, the flagellum is simple (i.e. neither dentate nor pectinate) but has its distal section somewhat sinuous and turned up apically (the original description of the Whalleyanidae (Minet 1991: 89) states “flagellum… on distal section curved as in the Callidulidae”; this antennal trait can also be seen in live adults (Wang 1993, photo of a Tetragonus catamitus Geyer, 1832), although often less distinctly; (8) on vertex, the chaetosemata are large and include minute scales between their setae; (9) veins Rs2 and Rs3 are stalked in the forewing (all Rs veins are “free” in many Thyrididae, *Prodidactis* (Janse 1964: pl. 5), *Hyblaea*, and in the ground plan of the Papilionoidea (cf. Hesperiidae)); (10) in the forewing, veins Rs3 and Rs4 run to the termen, reaching it below the apex (Minet 1998: Fig. 15.1, B and C; Viette 1977: Fig. 1) (by contrast, only Rs4 runs to the termen in *Prodidactis*, *Hyblaea*, and in the thyridid ground plan (cf. Common 1990: Fig. 109.4); nevertheless, through parallel evolution, several Thyrididae also possess apomorphy (10): (Common 1990:Fig. 109.1)); (11) in the basal region of the hindwing, there is a recurrent humeral spur, or fold (*Whalleyana*), between Sc and the frenulum (Minet 1998: Fig. 15.1, B and C); (12) in the hindwing, vein M2 arises much closer to M3 than to M1 (*Hyblaea* and most Thyrididae also have the hindwing vein M2 arising closer to M3 than to M1 but this vein arises midway between M1 and M3 in the thyridid ground plan (illustrated, for this character, by the genus Addaea Walker, 1866: Common 1990: Fig. 109.4) and arises slightly closer to M1 than to M3 in the hyblaeid genus *Erythrochrus* Herrich-Schäffer, 1858); (13) at the base of the abdomen, the marginotergites (term used by Brock (1971)) are anteriorly connected to the anterior angles of sternum A2 through complete tergosternal sclerites (Fig. 2B, long arrow) (in most Thyrididae, sternum A2 has just variously developed anterolateral processes that do not reach the marginotergites); (14) the apodemes of sternum A2 are short or reduced (Fig. 2B, short arrow) (although reduction of the apodemes also occurs in several Thyrididae, these structures are sometimes large or elongate in this family: e.g. Fig. 12 in Minet (1983)); (15) the male genitalia lack a complete gnathos (retained in many Thyrididae); (16) in the female genitalia, the eighth sternum is transversely elongate and distinctly arched (concave cephalad) (Fig. 2C; see also, for Callidulidae, several figures in Holloway (1998)). Among these ten derived traits, we regard (7), (8), (11), and (16) as really significant synapomorphies, which tend to support the results obtained with our molecular phylogenetic analysis. Therefore, we formally propose here to assign Whalleyanidae to the superfamily Calliduloidea (***revised concept***, with a definition based on apomorphies (7)-(16)).

Within the thus redefined Calliduloidea, nine autapomorphies support the monophyly of the family Callidulidae (= Pterothysaninae + Griveaudiinae + Callidulinae), namely: (17) foreleg with an apical pair of stronger spines on tarsomere 4, but with at most a few minute spines on the ventral surface of tarsomere 5 (Minet 1990: Figs 4-6); (18) male forewing without a subcostal retinaculum (while *Whalleyana vroni* retains this retinaculum (Fig. 2D, short arrow); through parallel evolution, the male forewing of *Whalleyana toni* has lost this structure); (19) in the forewing, anal vein simple, devoid of “basal fork” (A2 being at most a very short veinlet parallel to the base of vein A1 (Minet 1998: Fig. 15.1 B); by contrast, *Whalleyana* retains this “basal fork”, although with a weak lower branch: Fig. 2D, long arrow); (20) mesopleurosternum with the precoxal sulcus faintly indicated to wholly absent (unlike that observed in the two species of *Whalleyana*: Fig. 2E, arrow); (21) metascutellum less elongate, in posterior view (Fig. 2F, long arrow), than in *Whalleyana* (Fig. 2G, long arrow) and most moths; (22) fenestrae laterales very small (Fig. 2F, short arrow) (while they are well developed in both species of *Whalleyana* (Fig. 2G, short arrow) and rather large in most Thyrididae); (23) in the male genitalia, juxta without “erect arms” (a loss, as mentioned above: see (6)); (24) male genitalia with a short, sclerotized bridge, which unites the sacculi ventrad of the juxta (Minet 1990: Fig. 23); (25) female genitalia with a characteristic – flat and quadrilobate – ovipositor (see also Figs 21-25 in Holloway 1998; Minet 1990: Figs 27-29). Since Pterothysaninae and Griveaudiinae appear as sister groups on molecular evidence, it should be noted that the male genitalia also provide a synapomorphy for these two subfamilies, namely the presence of a few conspicuous setae in the membranous area situated just below the base of the uncus (Minet 1990:Figs 20 and 21).

## Discussion

### a) Molecular data

Here we present the results for low-coverage whole genome sequencing of Lepidoptera museum specimens and existing DNA extracts. With the exception of *H. puera* the DNA extracts used for library preparation were highly fragmented. *De novo* assembly resulted in highly fragmented assemblies (N50 range of 317 – 2,078 bp). These assemblies are consistent with assemblies obtained in other low-coverage whole genome sequencing projects, such as that of the swallowtail butterflies (Allio et al. 2020) and skipper butterflies (Li et al. 2019). Despite the highly fragmented nature of the resulting assemblies, the overall gene recovery rate was between 64% and 90%. Studies of bird museum specimens using AHE based approaches had recovery rates of 30 – 92% (Tsai et al. 2019) and 49 – 62% (McCormack et al. 2016). One advantage of sequencing genomes over target enrichment approaches, is that in the future one may go back to the original data or assembly and extract new sets of genes, rather than just being limited to the genes which were enriched for. The successful library preparation and high rate of gene recovery from the *H. puera* sample, highlights the usefulness of existing DNA extracts (which were originally extracted for PCR based studies) for whole genome sequencing. The ability to sequencing existing extracts that are sitting around in storage from previous studies represents an important resource for expanding not only our genetic datasets but our understanding of interesting taxa and questions which may have been previously limited due to the inability to collect fresh specimens for library construction.

We found that generating 10 times more sequence data for the *Whalleyana* specimen compared to the other four specimens did not lead to better *de novo* assembled genomes, or to higher recovery of gene regions of interest. It appears that approximately 20X coverage of a genome is enough to extract useful phylogenetic information from highly fragmented material. Lepidoptera tend to have fairly small genomes, approximately 500 Mb in size (Triant et al. 2018), thus making them amenable to pooling for sequencing on the Illumina platform, with about 10 genomes possible on a HiSeqX machine, or 60 genomes on the current NovaSeq machine.

We targeted 332 genes for our work, which were a set of genes that have been manually screened for orthology and alignment in a previous study (Rota et al. in prep). However, given that we have sequenced the whole genomes of our specimens, we would be able to bioinformatically extract much more information from them if necessary. Assessment of the genomes for the presence of single-copy core orthologues present in the insecta BUSCO lineage set, with a blast approach found the majority to genes were present, at least in a fragmented form. In our study, we are confident that the 332 genes have correctly placed *Whalleyana* in the Lepidoptera Tree of Life as sister to the family Callidulidae, and are reasonably confident that Hyblaeidae is sister to Pyraloidea.

### b) The morphological context

In our tree, a well-supported clade is composed of the Thyrididae, Whalleyanidae and Callidulidae (bootstrap support value: 100). The sister group to this clade is unclear, as discussed by Rota et al. (in prep.). Based on the amino acid dataset, Gelechioidea appear as the sister group of Thyrididae + Whalleyanidae + Callidulidae, or the latter is sister to Papilionoidea based on the nucleotide dataset. We did not find significant morphological evidence supporting either hypothesis (although Papilionoidea share three reductions/losses with Thyrididae + Whalleyanidae + Callidulidae, viz. the above-mentioned apomorphies (1), (3) and (4) (see section Results, b). All published phylogenomic analyses have been mainly based on amino acid data, and have placed Callidulidae and/or Thyrididae close to Gelechioidea (Bazinet et al., 2013; Kawahara & Breinholt, 2014; Kawahara et al., 2019). Given the instability of the relationship of the Callidulidae/Thyrididae clade, the above interpretation of morphological characters has taken into account the morphology of Gelechioidea and that of three other superfamilies, which have been associated with Thyrididae and/or Callidulidae in previous works (Mutanen et al., 2010; Kaila et al., 2013; Regier et al., 2013; Wahlberg et al., 2013; Heikkilä et al., 2015, etc.), namely the Papilionoidea, Hyblaeoidea and Pyraloidea.

The monotypic family Prodidactidae was convincingly assigned to the superfamily Hyblaeoidea by Kaila et al. (2013), notably on the basis of an unusual apomorphy found in the male hindcoxa (viz. a variously developed process arising from the coxal membrane and present in both *Prodidactis* and Hyblaeidae). These authors also found a previously unnoticed larval apomorphy (modified apex of the spinneret) in the two hyblaeoid families but also in the Thyrididae. Accordingly they regarded the spinneret modification as a possible synapomorphy of Hyblaeoidea and Thyridoidea. However they did not find clear molecular evidence supporting a sister-group relationship between these two superfamilies. It should be noted that Hyblaeidae and Thyrididae also share a possible forewing synapomorphy, namely a well defined bunch of piliform scales arising (dorsally) from the base of vein A1 (Fig. 2H, arrow). We found this apomorphic trait in the two hyblaeid genera (*Erythrochrus*; *Hyblaea*) and in all thyridid subfamilies but it does not exist in *Prodidactis* (Alma Solis, pers. comm.) so that it may correspond to a parallel evolution between Hyblaeidae and Thyrididae. Our molecular analysis tend to establish (like that of Heikkilä et al. 2015) a well supported sister-group relationship of Hyblaeoidea and Pyraloidea. Nevertheless we found only two possible synapomorphies for these superfamilies, namely the triangular shape, in lateral view, of the maxillary palps (due to the presence of a tuft of elongate scales: see e.g. Janse’s (1964) plate 21 for *Prodidactis*) and the closeness of the bases of M2 and M3 in the forewing venation (this apomorphy has probably arisen independently in Hyblaeoidea + Pyraloidea and Thyridoidea + Calliduloidea: cf. apomorphy (2)). The maxillary palp apomorphy may be significant: it occurs in several groups of Pyralidae (e.g. *Synaphe* Hübner, 1825) and Crambidae (Scopariinae, Heliothelinae, Crambinae, etc.) but must have been secondarily lost in many taxa (replaced with filiform or reduced maxillary palps).

### c) Conclusion

In conclusion, the results we present here show that good levels of gene recovery can be obtained from low-coverage whole genome sequencing of even highly fragmented museum samples. Our study highlights the usefulness of genome sequencing museum specimens for which we have very little prior knowledge, and lack the ability to collect fresh specimens. Additionally, we highlight that existing DNA extracts that were originally extracted for PCR are suitable for next-generation sequencing library preparation methods, and thereby represent a valuable untapped resource for expanding our datasets.

## Acknowledgements

We would like to thank: Nicolas Dussex for advice on library preparation protocols, Marko Mutanen for providing the *H. puera* DNA extract, and also Alma Solis and David C. Lees for helpful information (on *Prodidactis* and *Whalleyana* respectively). We acknowledge the support from the National Genomics Infrastructure in Genomics Production Stockholm funded by Science for Life Laboratory, the Knut and Alice Wallenberg Foundation and the Swedish Research Council, and SNIC/Uppsala Multidisciplinary Center for Advanced Computational Science for assistance with massively parallel sequencing and access to the UPPMAX computational infrastructure and the use of New Zealand eScience Infastructure (NeSI) high-performance computing facilities. New Zealand’s national facilities are provided by NeSI and funded jointly by NeSI’s collaborator institutions and through the Ministry of Business, Innovation & Employment’s Research Infrastructure programme. URL https://www.nesi.org.nz. This work was funded by the Swedish Research Council (grant no. 2015-04441)

## Notes

### Competing Interest Statement

The authors have declared no competing interest.

