## Supplementary Tables for "Museomics of a rare taxon: placing Whalleyanidae in the Lepidoptera Tree of Life"

**Table S1:** Library Preparation Reagents, Supplier and product code.

| Reagent or Resource | Source | Product Code |
|---|---|---|
| QIAamp DNA Micro Kit | Qiagen | 56304 |
| SYBR safe | Fisher Scientific | S33102 |
| 10X Buffer Tango | Fisher Scientific | 11541505 |
| dNTP | Sigma-Aldrich | DNTP100-KT |
| ATP | Sigma-Aldrich | 11140965001 |
| T4 DNA Polymerase | New England Biolab | M0203S |
| T4 Polynucleotide Kinase | New England Biolab | M0201S |
| Min Elute PCR purification Kit | Qiagen | 28006 |
| T4 DNA Ligase | New England Biolab | M0202 |
| PEG-4000 | Sigma-Aldrich | 1546569-1G |
| Bst 2.0 WarmStart DNA Polymerase | New England Biolab | M0538S |
| Accuprime Pfx DNA Polymerase | Fisher Scientific | 12344024 |
| Agencourt AMPure XP | Beckman Coulter | A63880 |
| Tween20 | Sigma-Aldrich | P9416 |
| EDTA | Fisher Scientific | 15575020 |
| Tris-Cl pH8 | Fisher Scientific | 10259194 |
| TE buffer | Sigma-Aldrich | 93283-100mL |
| NaCl solution | Sigma-Aldrich | 71386-1L |

**Table S2**: Complete list of taxa included in the analysis

| Code | Superfamily | Family | Genus | Species | NCBI accession | Type of data |
|---|---|---|---|---|---|---|
| NW_T143 | Bombycoidea | Apatelodidae | Apatelodes | pithala | SRR1794084 | Transcriptome |
| NW_G003 | Bombycoidea | Bombycidae | Bombyx | mori | GCF_000151625.1 | Genome |
| NW_T076 | Bombycoidea | Saturniidae | Actias | luna | SRR1002974 | Transcriptome |
| NW_T134 | Bombycoidea | Saturniidae | Antheraea | yamamai | SRR1743839 | Transcriptome |
| NW_T195 | Bombycoidea | Saturniidae | Attacus | atlas | SRR1002994 | Transcriptome |
| NW_T077 | Bombycoidea | Saturniidae | Eacles | imperialis | SRR1299435 | Transcriptome |
| NW_T131 | Bombycoidea | Saturniidae | Rhodinia | newara | SRR1743843 | Transcriptome |
| NW_T130 | Bombycoidea | Saturniidae | Samia | ricini | SRR1743841 | Transcriptome |
| NW_T078 | Bombycoidea | Saturniidae | Therinia | lactucina | SRR1299418 | Transcriptome |
| NW_T080 | Bombycoidea | Sphingidae | Adhemarius | daphne | SRR1299394 | Transcriptome |
| NW_T196 | Bombycoidea | Sphingidae | Ceratomia | undulosa | SRR1002985 | Transcriptome |
| NW_T197 | Bombycoidea | Sphingidae | Darapsa | myron | SRR1002986 | Transcriptome |
| NW_T081 | Bombycoidea | Sphingidae | Enyo | lugubris | SRR1002983 | Transcriptome |
| NW_T082 | Bombycoidea | Sphingidae | Hemaris | diffinis | SRR1002987 | Transcriptome |
| NW_G002 | Bombycoidea | Sphingidae | Manduca | sexta | GCF_000262585.1 | Genome |
| NW_T083 | Bombycoidea | Sphingidae | Xylophanes | tersa | SRR1298384 | Transcriptome |
| NW_T005 | Calliduloidea | Callidulidae | Pterodecta | felderi | SRR1299369 | Transcriptome |
| VT56 | Calliduloidea | Callidulidae | Helicomitra | pulchra | SRR11799524 | Genome |
| VT58 | Calliduloidea | Callidulidae | Griveaudia | vieui | SRR11799526 | Genome |
| NW_T108 | Carposinoidea | Carposinidae | Carposina | sasakii | GADL00000000 | Transcriptome |
| NW_T001 | Choreutoidea | Choreutidae | Hemerophila | diva | SRR1005851 | Transcriptome |
| NW_T006 | Cossoidea | Castniidae | Synemon | plana | SRR1006158 | Transcriptome |
| NW_T200 | Cossoidea | Castniidae | Telchin | licus | SRR1204999 | Transcriptome |
| NW_T049 | Cossoidea | Cossidae | Culama | ANIC3 | SRR1006166 | Transcriptome |
| NW_T222 | Cossoidea | Cossidae | Eogystia | hippophaecolus | SRR4409152 | Transcriptome |
| NW_T007 | Cossoidea | Cossidae | Givira | mucidus | SRR1006184 | Transcriptome |
| NW_T008 | Cossoidea | Cossidae | Morpheis | mathani | SRR1299214 | Transcriptome |
| NW_T050 | Cossoidea | Cossidae | Prionoxystus | robiniae | SRR1006187 | Transcriptome |

| | | | | | | |
|---|---|---|---|---|---|---|
| NW_T183 | Cossoidea | Cossidae | Psychogena | miranda | SRR1006219 | Transcriptome |
| NW_T052 | Cossoidea | Cossidae | Spinulata | maruga | SRR1006227 | Transcriptome |
| NW_T009 | Cossoidea | Dudgeoneidae | Archaeoses | polygrapha | SRR1021600 | Transcriptome |
| NW_T010 | Cossoidea | Sesiidae | Podosesia | syringae | SRR1021623 | Transcriptome |
| NW_T079 | Cossoidea | Sesiidae | Vitacea | polistiformis | SRR1021630 | Transcriptome |
| NW_T012 | Drepanoidea | Cimeliidae | Axia | margarita | SRR1006157 | Transcriptome |
| NW_T011 | Drepanoidea | Doidae | Doa | Janzen01 | SRR1021609 | Transcriptome |
| NW_T054 | Drepanoidea | Drepanidae | Cyclidia | substigmaria | SRR1021608 | Transcriptome |
| NW_T013 | Drepanoidea | Drepanidae | Pseudothyatira | cymatophoroides | SRR1021621 | Transcriptome |
| NW_T048 | Gelechioidea | Depressariidae | Psilocorsis | reflexella | SRR1021624 | Transcriptome |
| NW_T016 | Gelechioidea | Depressariidae | Tonica | nigricostella | SRR1299773 | Transcriptome |
| NW_T015 | Gelechioidea | Elachistidae | Antaeotricha | schlaegeri | SRR1021598 | Transcriptome |
| NW_T057 | Gelechioidea | Gelechiidae | Dichomeris | punctidiscella | SRR1021607 | Transcriptome |
| NW_T203 | Gelechioidea | Gelechiidae | Pectinophora | gossypiella | SRR2225787;<br>SRR2225788;<br>SRR2225790 | Transcriptome |
| NW_T125 | Gelechioidea | Gelechiidae | Tuta | absoluta | SRR1721964 | Transcriptome |
| NW_T066 | Gelechioidea | Lecithoceridae | Thubana | sp. | SRR1300991 | Transcriptome |
| NW_T014 | Geometroidea | Epicopeiidae | Epicopeia | hainseii | SRR1021610 | Transcriptome |
| NW_T017 | Geometroidea | Geometridae | Biston | betularia | SRR1021599 | Transcriptome |
| NW_T202 | Geometroidea | Geometridae | Biston | suppressaria | SRR1777716 | Transcriptome |
| NW_T058 | Geometroidea | Geometridae | Chlorosea | margaretaria | SRR1021603 | Transcriptome |
| NW_T153 | Geometroidea | Geometridae | Ectropis | obliqua | SRR3056076 | Transcriptome |
| NW_T059 | Geometroidea | Geometridae | Idaea | eremiata | SRR1021615 | Transcriptome |
| NW_T060 | Geometroidea | Geometridae | Macaria | distribuaria | SRR1299213 | Transcriptome |
| NW_T061 | Geometroidea | Geometridae | Nemoria | lixaria | SRR1299347 | Transcriptome |
| NW_T018 | Geometroidea | Sematuridae | Mania | lunus | SRR1299318 | Transcriptome |
| NW_T085 | Geometroidea | Uraniidae | Calledapteryx | dryopterata | SRR1021601 | Transcriptome |
| NW_T019 | Geometroidea | Uraniidae | Lyssa | zampa | SRR1299769 | Transcriptome |
| NW_T142 | Gracillarioidea | Gracillariidae | Acrocercops | transecta | DRR023321 | Transcriptome |
| NW_T146 | Gracillarioidea | Gracillariidae | Caloptilia | triadicae | SRR1794032 | Transcriptome |

| | | | | | | |
|---|---|---|---|---|---|---|
| NW_T087 | Gracillarioidea | Gracillariidae | Cameraria | ohridella | SRR1190496 | Genome |
| NW_T020 | Gracillarioidea | Gracillariidae | Phyllocnistis | citrella | SRR1299751 | Transcriptome |
| VT57 | Hyblaeoidea | Hyblaeidae | Hyblaea | madagascariensis | SRR11799525 | Genome |
| MM07227 | Hyblaeoidea | Hyblaeidae | Hyblaea | puera | SRR11799522 | Genome |
| NW_T023 | Immoidea | Immidae | Imma | tetrascia | SRR1021616 | Transcriptome |
| NW_T064 | Lasiocampoidea | Lasiocampidae | Artace | sp. | SRR1299316 | Transcriptome |
| NW_T151 | Lasiocampoidea | Lasiocampidae | Dendrolimus | houi | SRR944675 | Transcriptome |
| NW_T152 | Lasiocampoidea | Lasiocampidae | Dendrolimus | punctatus | SRR2641162 | Transcriptome |
| NW_T065 | Lasiocampoidea | Lasiocampidae | Tolype | notialis | SRR1021628 | Transcriptome |
| NW_T025 | Mimallonoidea | Mimallonidae | Lacosoma | chiridota | SRR1021619 | Transcriptome |
| NW_T024 | Mimallonoidea | Mimallonidae | Lacosoma | ludolpha | SRR1299212 | Transcriptome |
| NW_T144 | Noctuoidea | Erebidae | Arctia | plantaginis | ERR1856313 | Transcriptome |
| NW_T026 | Noctuoidea | Erebidae | Eudocima | salaminia | SRR1300148 | Transcriptome |
| NW_T156 | Noctuoidea | Erebidae | Euproctis | chrysorrhoea | SRR1040496 | Transcriptome |
| NW_T027 | Noctuoidea | Erebidae | Lymantria | dispar | SRR1021618 | Transcriptome |
| NW_T056 | Noctuoidea | Euteliidae | Anigraea | rubida | SRR1299755 | Transcriptome |
| NW_T106 | Noctuoidea | Noctuidae | Agrotis | segetum | GBCW00000000 | Transcriptome |
| NW_T107 | Noctuoidea | Noctuidae | Athetis | lepigone | GARD00000000 | Transcriptome |
| NW_T148 | Noctuoidea | Noctuidae | Chrysodeixis | includens | SRR2049082 | Transcriptome |
| NW_T069 | Noctuoidea | Noctuidae | Helicoverpa | zea | SRR1021614 | Transcriptome |
| NW_T163 | Noctuoidea | Noctuidae | Heliothis | subflexa | ERR738599 | Transcriptome |
| NW_T224 | Noctuoidea | Noctuidae | Mythimna | separata | SRR5115697 | Transcriptome |
| NW_T173 | Noctuoidea | Erebidae | Oraesia | emarginata | SRR5128005 | Transcriptome |
| NW_T187 | Noctuoidea | Noctuidae | Sesamia | nonagrioides | ERR424922 | Transcriptome |
| NW_T189 | Noctuoidea | Noctuidae | Spodoptera | exigua | SRR525279 | Transcriptome |
| NW_G008 | Noctuoidea | Noctuidae | Spodoptera | frugiperda | GCF_011064685.1 | Genome |
| NW_T071 | Noctuoidea | Noctuidae | Striacosta | albicosta | SRR037002 | Transcriptome |
| NW_T110 | Noctuoidea | Noctuidae | Trichoplusia | ni | GBKU00000000 | Transcriptome |
| NW_T072 | Noctuoidea | Nolidae | Manoba | major | SRR1300145 | Transcriptome |
| NW_T073 | Noctuoidea | Notodontidae | Notoplusia | minuta | SRR1299746 | Transcriptome |
| NW_T092 | Noctuoidea | Notodontidae | Thaumetopoea | pityocampa | SRR1284701 | Transcriptome |

| | | | | | | |
|---|---|---|---|---|---|---|
| NW_T029 | Papilionoidea | Hedylidae | Macrosoma | hedylaria | SRR1299306 | Transcriptome |
| NW_T116 | Papilionoidea | Hesperiidae | Erynnis | propertius | SRX016482 | Transcriptome |
| NW_T041 | Papilionoidea | Hesperiidae | Hylephila | phyleus | SRR1299296 | Transcriptome |
| NW_T062 | Papilionoidea | Hesperiidae | Megathymus | yuccae | SRR1299752 | Transcriptome |
| NW_T118 | Papilionoidea | Hesperiidae | Thymelicus | sylvestris | SRR1325130 | Transcriptome |
| NW_T193 | Papilionoidea | Hesperiidae | Urbanus | proteus | SRR1794082 | Transcriptome |
| NW_T068 | Papilionoidea | Lycaenidae | Hemiargus | ceraunus | SRR1299274 | Transcriptome |
| NW_T095 | Papilionoidea | Lycaenidae | Polyommatus | icarus | SRR921636 | Transcriptome |
| NW_T182 | Papilionoidea | Lycaenidae | Protantigius | superans | SRR2441452 | Transcriptome |
| NW_T188 | Papilionoidea | Lycaenidae | Spindasis | takanonis | SRR2441453 | Transcriptome |
| NW_G004 | Papilionoidea | Nymphalidae | Danaus | plexippus | GCA_000235995.2 | Genome |
| NW_T157 | Papilionoidea | Nymphalidae | Fabriciana | nerippe | SRR2837841 | Transcriptome |
| NW_G005 | Papilionoidea | Nymphalidae | Heliconius | melpomene | GCA_000313835.2 | Genome |
| NW_T088 | Papilionoidea | Nymphalidae | Junonia | coenia | SRR1166409 | Transcriptome |
| NW_T090 | Papilionoidea | Nymphalidae | Limenitis | arthemis | SRR1504907 | Transcriptome |
| NW_T171 | Papilionoidea | Nymphalidae | Maniola | jurtina | SRR3721684; SRR3721695 | Transcriptome |
| NW_G006 | Papilionoidea | Nymphalidae | Melitaea | cinxia | GCA_000716385.1 | Genome |
| NW_T089 | Papilionoidea | Nymphalidae | Pararge | aegeria | SRR1190479 | Genome |
| NW_T169 | Papilionoidea | Papilionidae | Luehdorfia | chinensis | SRR2924893 | Transcriptome |
| NW_T040 | Papilionoidea | Papilionidae | Papilio | glaucus | SRR850324 | Transcriptome |
| NW_T074 | Papilionoidea | Papilionidae | Papilio | polytes | SRR850327 | Transcriptome |
| NW_T098 | Papilionoidea | Papilionidae | Parides | eurimedes | SRR921629 | Transcriptome |
| NW_T123 | Papilionoidea | Pieridae | Anthocharis | cardamines | | Transcriptome |
| NW_T122 | Papilionoidea | Pieridae | Belenois | gidica | | Transcriptome |
| NW_T121 | Papilionoidea | Pieridae | Delias | nigrina | | Transcriptome |
| NW_T120 | Papilionoidea | Pieridae | Gonepteryx | rhamni | | Transcriptome |
| NW_T179 | Papilionoidea | Pieridae | Pieris | brassicae | ERR1855021 | Transcriptome |
| NW_T181 | Papilionoidea | Pieridae | Pontia | daplidice | SRR5253669 | Transcriptome |
| NW_T039 | Papilionoidea | Riodinidae | Semomesia | campanea | SRR1299211 | Transcriptome |
| NW_T003 | Pterophoroidea | Pterophoridae | Emmelina | monodactyla | SRR1021605 | Transcriptome |

| | | | | | | |
|---|---|---|---|---|---|---|
| NW_T002 | Pterophoroidea | Pterophoridae | Lantanophaga | pusillidactyla | SRR1299210 | Transcriptome |
| NW_T037 | Pyraloidea | Crambidae | Catoptria | oregonicus | SRR1021602 | Transcriptome |
| NW_G007 | Pyraloidea | Crambidae | Chilo | suppressalis | GCA_004000445.1 | Genome |
| NW_T053 | Pyraloidea | Crambidae | Cnaphalocrocis | medinalis | SRR647915 | Transcriptome |
| NW_T158 | Pyraloidea | Crambidae | Glyphodes | pyloalis | SRR5520571 | Transcriptome |
| NW_T159 | Pyraloidea | Crambidae | Haritalodes | derogata | SRR2017609 | Transcriptome |
| NW_T168 | Pyraloidea | Crambidae | Leucinodes | orbonalis | SRR5006982 | Transcriptome |
| NW_T223 | Pyraloidea | Crambidae | Loxostege | sticticalis | SRR4905822 | Transcriptome |
| NW_T172 | Pyraloidea | Crambidae | Maruca | vitrata | SRR074950 | Transcriptome |
| NW_T038 | Pyraloidea | Crambidae | Myelobia | smerintha | SRR1299267 | Transcriptome |
| NW_T109 | Pyraloidea | Crambidae | Ostrinia | nubilalis | GAVD00000000 | Transcriptome |
| NW_T185 | Pyraloidea | Crambidae | Scirpophaga | incertulas | SRR1615983; SRR1615982; SRR1613323 | Transcriptome |
| NW_T190 | Pyraloidea | Crambidae | Spoladea | recurvalis | SRR1511612 | Transcriptome |
| NW_T201 | Pyraloidea | Pyralidae | Cadra | cautella | SRR1508178 | Transcriptome |
| NW_T075 | Pyraloidea | Pyralidae | Galleria | melonella | SRR1021612 | Transcriptome |
| NW_T114 | Pyraloidea | Pyralidae | Plodia | interpunctella | ERR426341 | Transcriptome |
| NW_T084 | Thyridoidea | Thyrididae | Pseudothyris | sepulchralis | SRR1299495 | Transcriptome |
| NW_T042 | Thyridoidea | Thyrididae | Striglina | suzukii | SRR1021625 | Transcriptome |
| NW_T043 | Thyridoidea | Thyrididae | Zeuzerodes | maculata | SRR1299209 | Transcriptome |
| NW_T044 | Tineoidea | Psychidae | Thyridopteryx | ephemeraeformis | SRR1021627 | Transcriptome |
| NW_T209 | Tineoidea | Dryadaulidae | Dryadaula | visaliella | SRR3180631 | Transcriptome |
| NW_T211 | Tineoidea | Meessiidae | Eudarcia | simulatricella | SRR3180633 | Transcriptome |
| NW_T216 | Tineoidea | Tineidae | Tineola | bisselliella | SRR3180622 | Transcriptome |
| NW_T149 | Tortricoidea | Tortricidae | Ctenopseustis | herana | SRR1165709 | Transcriptome |
| NW_T045 | Tortricoidea | Tortricidae | Cydia | pomonella | SRR1021606 | Transcriptome |
| NW_T137 | Tortricoidea | Tortricidae | Epiphyas | postvittana | SRR1151338 | Transcriptome |
| NW_T111 | Tortricoidea | Tortricidae | Grapholita | molesta | GADK00000000 | Transcriptome |
| NW_T160 | Tortricoidea | Tortricidae | Hedya | nubiferana | SRR3476345 | Transcriptome |
| NW_T047 | Tortricoidea | Tortricidae | Phaecasiophora | niveiguttana | SRR1021604 | Transcriptome |

| | | | | | | |
|---|---|---|---|---|---|---|
| NW_T113 | Tortricoidea | Tortricidae | Planotortrix | excessana | SRR1258863 | Transcriptome |
| NW_T115 | Tortricoidea | Tortricidae | Rhyacionia | leptotubula | DRR012970 | Transcriptome |
| NW_T030 | Urodoidea | Urodidae | Urodus | decens | SRR1021629 | Transcriptome |
| NW_T031 | Urodoidea | Urodidae | Urodus | parvula | SRR1299750 | Transcriptome |
| VT11 | Whalleyanoidea | Whalleyanidae | Whalleyana | vroni | SRR11799523 | Genome |
| NW_G001 | Yponomeutoidea | Plutellidae | Plutella | xylostella | GCF_000330985.1 | Genome |
| NW_T097 | Yponomeutoidea | Yponomeutidae | Yponomeuta | evonymella | SRR921659 | Transcriptome |
| NW_T033 | Zygaenoidea | Dalceridae | Dalcera | abrasa | SRR1299208 | Transcriptome |
| NW_T055 | Zygaenoidea | Epipyropidae | Epipomponia | nawai | SRR1021611 | Transcriptome |
| NW_T063 | Zygaenoidea | Lacturidae | Lactura | subfervens | SRR1021626 | Transcriptome |
| NW_T067 | Zygaenoidea | Limacodidae | Euclea | delphinii | SRR1021597 | Transcriptome |
| NW_T034 | Zygaenoidea | Megalopygidae | Megalopyge | crispata | SRR1021617 | Transcriptome |
| NW_T035 | Zygaenoidea | Megalopygidae | Megalopyge | tharops | SRR1299217 | Transcriptome |
| NW_T036 | Zygaenoidea | Zygaenidae | Zygaena | fausta | SRR1021632 | Transcriptome |
| NW_T194 | Zygaenoidea | Zygaenidae | Zygaena | filipendulae | SRR023844 | Transcriptome |

**Table S3**: Complete summary of 332 gene recovery for specimens sequenced in this study. Presence of the gene in the dataset is represented by an X. Gene codes corresponds to the codes used with Rota et al (in prep).

| Gene | VT56 | VT58 | VT57 | MM07227 | VT11 |
|---|---|---|---|---|---|
| 14-3-3E | - | - | - | - | X |
| 2ODHa | X | X | X | - | - |
| 2ODHb | X | - | - | X | X |
| 39SrpL24 | X | - | X | X | X |
| 6PFK | X | - | X | X | X |
| 6PGD | - | X | X | X | X |
| ACC | X | X | X | - | - |
| Actin2 | - | - | X | X | - |
| acyl | - | X | - | X | X |
| ADF1 | X | - | X | X | X |
| ADH | X | - | X | X | X |
| AdK2 | X | - | X | X | X |
| ADP-RF1 | - | - | - | - | - |
| ADPGK | - | X | X | X | X |
| ADPRF | X | X | X | X | X |
| AFG3 | X | X | - | X | X |
| AGBE | X | X | X | X | X |
| AH | X | X | X | X | - |
| AIa | X | X | - | X | X |
| AIb | X | - | X | X | X |
| AK3 | X | X | X | X | X |
| Ala | X | X | - | X | - |
| AlDH | - | X | - | X | X |
| ALG2 | X | - | - | X | - |
| ANK13C | X | X | - | X | X |
| ArgKin | X | X | X | X | - |
| ATP6 | X | X | X | X | X |
| ATPase1 | - | - | - | X | - |
| ATPase6 | - | - | X | X | X |
| ATPasea | X | X | - | X | X |
| ATPaseAC39 | X | X | X | X | X |
| ATPaseb | X | X | - | X | X |
| ATPaseC | - | X | X | X | X |
| ATPaseDelta | X | X | X | X | X |
| ATPasee | X | - | X | X | - |
| ATPaseEpsil | X | X | - | - | - |
| ATPaseF | X | X | X | X | X |

| Gene | VT56 | VT58 | VT57 | MM07227 | VT11 |
|---|---|---|---|---|---|
| PDHE1 | X | X | X | X | X |
| PG | X | X | - | X | X |
| PGK | X | - | X | X | - |
| PGLYM | X | X | X | X | - |
| PIXFT | - | X | X | - | X |
| PK | X | X | X | X | X |
| Pleck | X | X | - | X | X |
| PolII | X | X | X | X | X |
| Porin | X | - | X | X | X |
| ProSup | X | X | X | X | - |
| PS | - | X | X | X | X |
| PSb | - | X | X | X | X |
| R5PI | X | X | - | X | - |
| RAB5 | X | - | - | X | X |
| Ras | X | X | X | X | X |
| RGNE | X | - | X | X | X |
| RP3E | - | - | - | - | - |
| RPF2 | - | - | - | X | X |
| RpL10 | X | X | X | X | X |
| RpL10A | X | X | X | X | X |
| RpL11 | X | X | X | X | X |
| RpL12 | X | X | X | X | X |
| RpL13 | X | X | - | X | X |
| RpL13A | X | X | - | X | X |
| RpL14 | - | - | X | X | X |
| RpL15 | X | X | X | X | X |
| RpL17 | X | X | X | X | X |
| RpL18 | X | X | X | X | X |
| RpL18A | - | X | X | X | - |
| RpL19 | - | - | X | X | - |
| RpL2 | X | - | X | X | X |
| RpL20 | X | X | X | X | X |
| RpL21 | X | - | - | X | X |
| RpL22 | X | - | X | X | X |
| RpL23 | X | X | X | X | X |
| RpL24 | X | X | X | X | X |
| RpL24A | - | X | X | X | X |

| | | | | | |
|---|---|---|---|---|---|
| ATPaseg | X | - | X | X | - |
| ATPaseH | X | X | X | - | - |
| ATPaseIa | - | - | X | X | - |
| ATPaseIb | - | - | - | X | - |
| ATPaseIc | - | - | - | X | - |
| ATPaseOS | X | - | X | X | X |
| ATPaseProt | X | X | X | X | X |
| ATPasesubD | X | - | X | X | X |
| ATPCL | X | X | X | - | X |
| ATPgamma | X | X | X | X | X |
| ATreP | - | X | X | X | - |
| BPx1 | X | X | X | X | X |
| BTB | X | - | X | X | X |
| Ca-ATPase | - | - | - | - | - |
| Ca2 | - | X | - | X | X |
| CAD | X | X | X | X | X |
| CAH | - | X | X | X | X |
| CDK5 | - | - | X | X | - |
| Chap6a | X | X | X | X | X |
| CHIP | - | - | - | X | X |
| chitinase | X | X | - | X | X |
| CHL | - | X | X | X | X |
| CMBP5 | X | X | X | X | X |
| COCC | X | X | - | X | X |
| COI | X | X | X | X | X |
| COII | X | X | X | X | X |
| COIII | X | X | X | X | X |
| COIV | X | X | X | X | X |
| COP9 | X | X | X | - | X |
| COVa | X | X | X | X | X |
| COVb | - | X | X | - | X |
| COVIA1 | X | - | - | X | X |
| COVIIc1 | - | - | X | X | - |
| COX11 | X | X | X | X | X |
| CRc1 | - | X | X | X | X |
| CRis | - | X | X | X | X |
| CS | - | X | X | X | - |
| Cullin5 | X | X | X | X | X |
| CycH | X | - | X | X | X |
| CycY | - | X | X | X | X |

| | | | | | |
|---|---|---|---|---|---|
| RpL26 | X | - | X | X | X |
| RpL27 | - | X | X | X | X |
| RpL27A | - | - | X | X | - |
| RpL28 | X | X | X | X | X |
| RpL29 | X | X | - | X | - |
| RpL3 | - | - | X | X | X |
| RpL30 | X | - | X | X | X |
| RpL31 | - | X | - | X | X |
| RpL32 | X | X | X | - | X |
| RpL34 | - | X | X | X | X |
| RpL35 | X | - | X | - | X |
| RpL35A | X | X | X | X | - |
| RpL36 | - | - | X | X | - |
| RpL36A | X | - | X | X | X |
| RpL37 | X | X | - | X | X |
| RpL37A | - | X | X | X | X |
| RpL38 | X | X | X | X | - |
| RpL39 | X | - | X | X | - |
| RpL4 | - | X | - | X | X |
| RpL5 | X | X | X | X | - |
| RpL7 | X | - | X | X | X |
| RpL7A | X | - | X | X | X |
| RpL8 | - | - | X | X | X |
| RpL9 | X | X | X | X | X |
| RpP0 | X | - | X | X | X |
| RpP1 | X | - | X | X | X |
| RpP2 | - | - | X | X | - |
| RpS10 | - | X | X | X | X |
| RpS11 | X | X | - | X | X |
| RpS12 | - | - | X | X | X |
| RpS13 | X | X | X | X | X |
| RpS14 | X | X | X | X | - |
| RpS15 | X | X | X | X | X |
| RpS15A | X | - | X | X | X |
| RpS16 | X | - | - | X | X |
| RpS17 | X | X | X | X | X |
| RpS18 | X | X | - | X | - |
| RpS19 | X | - | - | X | X |
| RpS2 | - | - | X | X | X |
| RpS20 | X | - | X | X | X |

| | | | | | |
|---|---|---|---|---|---|
| CysP | X | X | X | X | - |
| CytB | X | X | X | X | X |
| CytB9 | X | X | - | - | X |
| CytBc18 | - | - | X | X | - |
| DDB1 | - | X | X | X | X |
| DDC | X | - | - | X | X |
| DDPS | X | X | X | X | X |
| DDX10 | X | X | - | X | - |
| DDX23 | - | X | X | X | - |
| DLDH | X | - | X | X | - |
| DnaJ | X | X | X | X | X |
| E2C | - | - | - | X | X |
| EAP30 | - | X | X | X | - |
| EF1a | X | - | - | X | X |
| EF1gA | - | X | - | X | - |
| EF1gB | X | - | - | - | - |
| eno | X | - | - | X | - |
| Exp1 | - | X | X | X | X |
| F16BA | X | - | - | X | - |
| F16BP | X | - | X | X | - |
| FCF1 | X | X | X | X | X |
| FH | X | X | X | X | X |
| ForKin | X | X | X | - | - |
| G1PU | - | X | - | X | X |
| G6P1E | X | - | X | X | X |
| GAPDH | X | - | X | X | X |
| gels | X | - | X | X | X |
| GLYP | X | X | X | X | X |
| GPD | X | - | - | X | X |
| GPI | - | X | X | X | - |
| gpts | X | - | X | X | X |
| GSS | X | - | X | - | X |
| GTPb | X | X | X | X | X |
| GTPCHI | - | X | X | X | X |
| HBS1 | X | - | X | X | X |
| HCADH | X | - | - | X | - |
| his | X | X | X | X | X |
| HK | - | - | X | X | - |
| IAP | X | - | X | X | - |
| IDH | X | X | X | X | X |

| | | | | | |
|---|---|---|---|---|---|
| RpS21 | X | X | X | X | - |
| RpS23 | X | X | X | X | - |
| RpS24 | - | X | X | X | X |
| RpS25 | - | X | X | X | X |
| RpS26 | X | X | - | X | X |
| RpS27 | X | - | X | X | X |
| RpS27A | - | X | - | - | X |
| RpS28 | - | - | X | X | X |
| RpS29 | X | X | - | X | X |
| RpS2b | X | X | - | X | X |
| RpS3 | X | X | X | X | X |
| RpS30 | - | - | - | X | X |
| RpS3A | - | X | X | X | X |
| RpS4 | X | - | - | - | X |
| RpS5 | X | X | X | X | X |
| RpS6 | - | X | - | X | X |
| RpS7 | X | - | X | X | X |
| RpS8 | X | X | X | X | X |
| RpS9 | X | - | X | X | X |
| RpSA | X | X | X | X | - |
| RSP1 | X | X | X | X | - |
| SARAH | X | X | X | X | X |
| SDH | X | X | X | X | - |
| SDHFS | X | X | X | X | X |
| SF38A | - | X | - | X | X |
| SL | X | X | X | X | X |
| slowmo | X | X | X | - | X |
| SOD | - | X | X | X | X |
| SPT16 | - | - | X | X | - |
| Ssu72 | X | - | - | X | X |
| STPP | X | X | - | X | - |
| SucCL | X | X | X | X | X |
| SucCoA | - | X | X | X | X |
| SucCoAb | X | X | X | X | - |
| TA | X | - | X | - | X |
| TBP | X | X | - | X | X |
| TfB | - | - | - | X | X |
| TFIIF | X | X | X | X | X |
| TFS | X | - | X | X | X |
| TGF | X | - | X | X | X |

| | | | | | |
|---|---|---|---|---|---|
| IDHa | X | X | - | X | X |
| IDHb | X | X | - | X | X |
| IDHc | X | X | X | X | X |
| IEBF | X | X | X | - | X |
| IF5C | X | - | X | X | X |
| JHBP | X | X | X | X | X |
| JM | X | X | X | - | X |
| KRR1 | X | - | - | X | X |
| LDH | - | - | X | - | X |
| LeuZip | X | X | - | X | X |
| LIS | - | - | X | X | - |
| LPL | - | - | X | X | X |
| luc7 | X | X | X | - | - |
| M1PD | X | X | X | X | X |
| MalDH | X | - | - | X | X |
| MaNa | X | X | X | X | X |
| MDH | X | X | X | X | X |
| MedPolII | X | X | X | X | X |
| MK6 | X | X | X | X | X |
| MMP41 | X | X | - | X | X |
| MPP2 | X | X | X | X | X |
| NAT10 | X | X | - | X | X |
| NC | X | X | X | X | X |
| ND1 | X | X | X | X | X |
| ND1a1 | X | X | X | - | X |
| ND1a10 | X | X | X | - | X |
| ND1a12 | X | - | - | X | - |
| ND1a13 | X | X | - | X | X |
| ND1a2 | - | X | - | X | X |
| ND1a4 | - | - | X | X | - |
| ND1a5 | - | - | X | X | X |
| ND1a6 | X | X | X | X | X |
| ND1a7 | - | X | - | - | X |
| ND1a8 | - | X | X | - | - |
| ND1a9 | X | X | X | X | - |
| ND1ab1 | X | X | - | X | X |
| ND1b1 | X | - | - | X | X |
| ND1b10 | X | X | - | X | - |
| ND1b5 | X | - | X | X | X |
| ND1b6 | - | X | - | X | X |

| | | | | | |
|---|---|---|---|---|---|
| TIF2A | X | X | X | X | X |
| TIF3B | X | X | X | X | X |
| TIF3Ca | X | X | - | X | X |
| TIF3Cb | - | X | X | X | X |
| TIF3K | X | X | X | X | X |
| TIF3M | X | X | X | X | - |
| TIF4A | X | - | X | X | X |
| TIF5A | - | - | X | X | - |
| TIF6 | - | X | X | X | X |
| TK | X | - | X | - | - |
| TP50A | X | - | X | X | X |
| TPI | X | X | X | X | X |
| TPPC11 | - | - | X | X | X |
| Treh-2 | - | - | - | - | - |
| Trop | X | X | X | X | X |
| TRP | - | - | - | X | - |
| U1A | X | X | X | X | X |
| U5 | - | X | X | X | - |
| UBCc | X | X | X | X | X |
| UCCR7 | X | X | X | X | - |
| UCTH | X | X | - | X | X |
| UDPG6DH | X | X | X | X | X |
| UDPGAD | X | X | X | X | X |
| UnP | X | X | X | X | - |
| vacA | X | X | - | X | X |
| vacATPc | - | X | X | X | - |
| vacATPd | - | X | X | X | X |
| vacATPE | X | X | X | X | X |
| vacATPF | X | - | X | X | X |
| vacATPg | X | X | X | X | X |
| vacATPH | X | X | X | X | X |
| vacB | X | - | - | X | - |
| VATPc | X | - | - | X | X |
| VPS26 | X | X | X | X | X |
| VPS4 | - | X | X | X | X |
| WD40 | X | X | X | X | X |
| WPH | X | X | - | X | X |
| ZN330 | X | - | - | X | X |

| | | | | | |
|---|---|---|---|---|---|
| ND1b9 | X | - | X | X | - |
| ND2 | X | - | X | X | X |
| ND3 | X | X | X | X | X |
| ND4 | X | X | X | X | X |
| ND4L | X | X | X | X | X |
| ND5 | X | X | X | X | X |
| NDF1 | X | X | X | X | X |
| NDF2 | X | X | - | X | X |
| NDFS1 | - | - | X | - | - |
| NDFS2 | X | X | - | X | X |
| NDFS3 | X | - | - | X | X |
| NDFS7 | X | X | X | X | X |
| NDFS8 | X | X | - | X | X |
| NEDD8 | X | X | X | X | X |
| Nex9 | X | X | X | X | X |
| nGTPb1 | X | - | X | X | X |
| NIDH | X | - | X | X | X |
| NIF3 | X | X | - | X | X |
| NMD3 | - | - | X | X | X |
| nuc56 | X | X | X | X | X |
| nuc58 | - | X | - | X | - |
| OSGEP | X | - | X | X | X |
| P26S12 | X | - | - | X | X |
| P26S4 | X | - | X | X | X |
| PCNA | - | - | X | X | X |
| PDH | X | - | X | - | - |
